# Fitting and Selection of Within-Host Models for SARS-CoV-2 Infection

**DOI:** 10.64898/2026.10.15.759195

**Authors:** N. Juhász, D. Selěsi, D. Nagy, B. Das, F. A. Bartha, D. Rajter-Ćirić, G. Röst

**Author notes:** Corresponding author: Nóra Juhász.

## Abstract

The main question in model selection for longitudinal RT-qPCR data is whether improvements in fit justify the additional parameters of more complex within-host models. We compared a target-cell-limited model (TIV), an eclipse-phase extension (TEIV), and a fractional-order TIV using infection waves reconstructed from SARS-CoV-2 cycle-threshold (Ct) measurements in a National Basketball Association cohort. Each trajectory was fitted on the Ct scale by maximizing a right-censored Gaussian likelihood using hybrid global–local optimization. Model selection was based primarily on the small-sample corrected Akaike information criterion (AICc), computed from the censored likelihood; while robustness to the choice of information criterion was assessed using the Akaike information criterion (AIC) and Bayesian information criterion (BIC). Among 868 evaluable trajectories, the unpenalized likelihood ranking identified TEIV as the clear best-fitting model in 34.0% of cases, fr-TIV in 30.2%, and TIV in 3.3%; the remaining 32.5% were effectively tied. After accounting for model complexity, AICc clearly favored TIV in 97.8% of trajectories and fr-TIV in 0.5%, while 1.7% were effectively tied; TEIV was not a clear winner in any of the trajectories. Thus, for these short, sparsely sampled Ct-only trajectories, TIV provided the best-supported parsimonious effective description. This does not imply that eclipse-phase or memory-like dynamics are biologically absent, but rather that the simpler TIV model is favoured by standard information criteria for the limited information contained in these data.

## 1 Introduction

Within-host mathematical models constitute a widely used framework for describing viral dynamics, with origins in foundational studies of HIV infection [1, 2]. They have subsequently been applied to estimate parameters governing viral kinetics from longitudinal or experimental time-course data across a broad range of viral infections [3, 4], including SARS-CoV-2 [5, 6, 7, 8, 9]. Despite their broad application, these models need not provide a unique mechanistic interpretation of longitudinal viral-load data: competing within-host models can produce similar viral-RNA trajectories while attributing viral proliferation and clearance to different mechanisms. This problem is particularly relevant when only longitudinal RT-qPCR measurements are available, because such data may support relatively simple target-cell-limited descriptions without separately identifying eclipse-phase, immune-response, or other extensions [10, 11]. Model comparison must therefore ask not only which formulation provides the best fit to the observations, but whether any improvement is significant enough to justify the additional parameters arising from more complex models.

The NBA occupational health program generated a large longitudinal dataset through repeated SARS-CoV-2 RT-qPCR testing of individuals affiliated with the league between July 2020 and January 2022 [12]. The resulting within-person Ct series follow individual acute infections and provide a common observational basis for comparing alternative within-host models. In this study, we use these trajectories to evaluate model support at the level of individual infection waves. Data from the program have previously been analyzed for related but distinct purposes. Kissler et al. fitted a Bayesian hierarchical, piecewise-linear model to Ct-derived viral concentrations [13]. Hay et al. subsequently analyzed 1,280 infections to investigate the effects of immune history, viral variant, and age [12]. These studies characterized viral trajectories, but did not compare mechanistic target-cell models. By contrast, Ke et al. used trajectories from nine intensively sampled NBA individuals, together with an independent German dataset, for an AIC-based comparison using a nonlinear mixed-effects approach [14].

Related model comparisons have been performed for other SARS-CoV-2 datasets as well, although the scope and design of these studies differed from ours. In particular, a comparison of mechanistic SARS-CoV-2 within-host models was performed by Hernandez et al. using viral-load measurements from nine patients [10]. Their fit included a target-cell-limited model and an extension with an eclipse phase; the models were compared using the corrected Akaike information criterion. In their analysis, the eclipse-phase extension was not supported by the available observations. They also emphasized the identifiability limitations that arise when viral load is the only measured model output. Ke et al. subsequently compared five mechanistic within-host models using daily RT-qPCR measurements from 56 participants in an independent cohort [15]. These works provide important precedents for information-criterion-based selection among SARS-CoV-2 models, while leaving open how such comparisons behave for different models, and generalize to larger collections of longitudinal data.

The relationship between model complexity and the information content of viral-load data was examined further by Sachak-Patwa et al. in the context of influenza infection [11]. They compared a hierarchy consisting of an explicit immune-response model, a TEIV model, and a simpler target-cell-limited formulation. When immune-response and cell-level measurements were available alongside viral load, the parameters of the immune-response model could be identified. With viral-load data alone, however, the simpler models were better supported.

In addition to the classical TIV and TEIV formulations considered in earlier studies, our comparison includes a fractional-order extension of the TIV model as a novelty. Fractional-order formulations provide another way of extending classical within-host models. Rather than introducing an additional biological compartment, they allow the present dynamics to depend on the previous trajectory of the system through a fractional-order parameter [16]. This additional degree of freedom may improve agreement with observed trajectories, but information-criterion-based comparisons weigh such improvements against the accompanying increase in model complexity. Previous analyses of the NBA data did not examine whether this history-dependent extension is supported when compared with classical TIV and TEIV models using information criteria.

In this study, we reconstruct individual infection waves from SARS-CoV-2 Ct measurements reported by Hay et al. [12] and fit three candidate models separately to each trajectory: a target-cell-limited TIV model, a TEIV model containing an eclipse phase, and a fractional-order TIV model. Because the TEIV and fractional-order models introduce additional parameters, improved goodness of fit alone is not sufficient to establish that they provide a better description of the data. We therefore distinguish unpenalized fit from complexity-penalized model selection by evaluating negative log-likelihood separately from AICc, with AIC and BIC used as sensitivity analyses. We also examine the sensitivity of the comparison to the Ct-to-viral-load calibration. Our aim is to assess whether the available Ct-only trajectories contain sufficient information to support additional parameterization of eclipse-phase or history-dependent dynamics.

## 2 Methods

### 2.1 Data source and wave extraction

We use longitudinal SARS-CoV-2 RT-qPCR testing data from the US National Basketball Association (NBA) cohort, collected between July 7, 2020 and January 26, 2022 [12]. Clinical specimens were collected from the anterior nares and oropharynx.

We defined individual infection waves as chronologically ordered sequences of Ct measurements for one participant during a single recorded infection episode. A wave was considered to end once at least three consecutive Ct measurements of 40 (i.e., no detectable viral RNA) followed an earlier within-episode detection, marking sustained clearance; isolated near-threshold measurements occurring later were treated as post-clearance noise rather than part of the acute viral trajectory and were excluded from the fitted wave.

For model fitting, we further retained only waves with at least seven uncensored Ct measurements (*Ct <* 40) and at least one Ct value at or below 35. The seven-point threshold was chosen to provide more uncensored observations than the maximum number of estimated parameters (six). The *Ct ≤* 35 requirement ensured that each fitted trajectory captured genuine evidence of an acute viral-load episode rather than only near-threshold detections. We additionally discarded waves with fewer than 8 time points to satisfy the AICc requirement *n > k* + 1, where *n* is the number of observations and *k* is the number of estimated parameters in a given model. The most complex models in this study had five fitted parameters, including the time offset, plus one estimated residual-variance parameter, giving *k* = 6 and thus requiring *n ≥* 8. Applying these criteria retained 868 of the original 1,280 waves for model fitting.

### 2.2 Ct measurements and viral-load calibration

Viral RNA in the NBA cohort was quantified by RT-qPCR using the Roche cobas SARS-CoV-2 assay [12], which reports a cycle threshold (Ct) for Target 1 (in this case, ORF1 a/b region): the number of amplification cycles required for the fluorescent signal to cross a fixed detection threshold. Since each cycle approximately doubles the amplified signal, Ct is inversely related to viral RNA concentration.

Ct-to-viral-load calibrations are assay- and target-gene-specific rather than universal: different PCR platforms and probe targets yield different Ct-to-copy-number relationships for the same underlying viral concentration [17, 13]. Because the NBA cohort was tested with the Roche cobas Target 1 assay, for which no dedicated copy-number standard curve is available, we used the two-step calibration of Kissler et al. [13, 12]: Target 1 Ct values were first cross-calibrated against Ct values from a validated CDC N1 assay by linear regression on paired specimens,

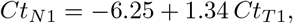

and the N1 Ct was then converted to genome copy number using a standard curve built from serial dilutions of a synthetic RNA transcript,

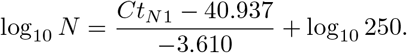

Composing these two steps gives a direct relationship between the observed Target 1 Ct and viral concentration *v* (copies/mL):

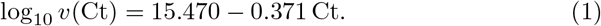

In this study, the models are formulated in terms of viral load *V* (*t*), whereas the directly observed measurements are Ct values. We therefore used the calibration curve as the observation link between the model state and the data. Let *V_M_* (*t*; ***θ****_M_*) denote the viral-load trajectory predicted by model *M*, where ***θ****_M_* contains its fitted dynamical parameters. Then, for model *M*, the untruncated Ct-scale mean (measurement noise will be discussed later) corresponding to the simulated viral load at sampling time *x_i_*, after applying the time offset *τ*, is

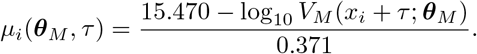

This latent mean was not truncated at Ct = 40 when evaluating the likelihood. Consequently, model predictions corresponding to viral loads below *v*(40) could have *µ_i_ >* 40, preserving the distinction between different viral loads below the reporting threshold.

### 2.3 Candidate within-host models

We consider three within-host models of SARS-CoV-2 infection: the targetcell-limited TIV model, its eclipse-phase extension (TEIV), and a fractional-order extension of TIV. The candidate set was designed to test whether the data support moving beyond the TIV baseline, using TEIV and fr-TIV as two prominent one-parameter extensions. All three models describe the same core infection process: target cells become infected, infected cells produce virus, infected cells die, and free virus is cleared. They differ in how this basic structure is extended: TEIV introduces an explicit eclipse phase, whereas the fractional-order model incorporates history-dependent dynamics without adding a new biological compartment.

The implemented candidate models are compared as distinct model families rather than as a nested sequence, so the profiled negative log-likelihood is not guaranteed to decrease as model complexity increases. In particular, the fractional-order model excludes the endpoint where it would coincide with TIV; while the fitted TEIV model reduces to TIV only in the limit of an infinitely fast eclipse-phase transition, which lies outside the finite parameter range used in our fits.

Following the measurement-scale convention used by [10], in all three models *V* is interpreted as RNA copies per mL on the RT-qPCR swab-sample scale. We highlight that it is not the local concentration of infectious virus at the target-cell surface in the airway. Consequently, fitted model parameters should generally be read as effective coefficients calibrated to sampling, dilution, and assay scale, not as pure physiological constants. For example, if *p*_airway_ denotes the true per-cell viral production rate in the infected airway and *q* denotes the combined recovery, transport, dilution, and assay-sampling factor linking airway virus to measured swab RNA copies, then the production coefficient on the observed RNA-copy scale is an effective quantity that includes *q · p*_airway_, similarly as in other swab-scale SARS-CoV-2 viral-dynamics models [18].

In all three models, the cellular state variables are normalized to the initial target-cell level, with *T*_0_ = 1. Moreover, in all models, *D* is an auxiliary state that records cumulative infected-cell death and does not feed back into the remaining dynamics.

None of the three candidate models described below includes an explicit immune-response component. This choice is motivated by the type of data available. Sachak-Patwa et al. found that the parameters of an explicit immuneresponse model could be identified when the model was fitted to data that jointly included information on viral-load and immune-response. When only viral-load data were used, however, they could estimate parameters of the TEIV and TIV models but not a more complex model incorporating an explicit immune-response mechanism [11]. Because our dataset likewise contains only longitudinal Ct measurements, we restrict the candidate set to models without explicit immune-response compartments. Immune effects may nevertheless be reflected implicitly in effective parameters such as the infected-cell death rate *δ* and the viral clearance rate *c*.

Because the analyzed SARS-CoV-2 infections evolve over approximately one to two weeks, a timescale much shorter than the turnover of uninfected respiratory epithelial cells, target-cell production and natural death are neglected in all models.

#### 2.3.1 TIV model

The TIV model [11] is the simplest candidate and serves as the reference formulation for the model comparison. It is defined by

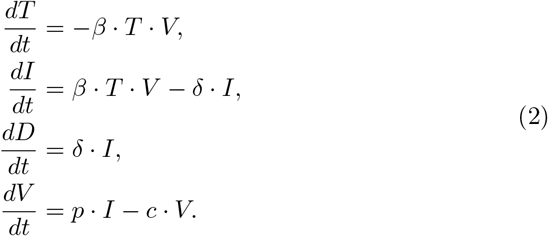

In system (2), *T* denotes uninfected target cells, *I* infected cells, *D* dead infected cells, and *V* the viral RNA concentration.

We use the TIV formulation of [10], with *V* and the associated rate parameters interpreted on the swab-measurement scale as described above.

In the TIV model, target cells become infected through mass-action kinetics at rate *βTV* [19]. Infected cells produce virus at rate *p* and die at rate *δ*, while free virus is cleared at rate *c*.

The structure of the TIV model is illustrated in panel A in Figure 1, while the model parameters are summarized in Table 1.

**Figure 1:**
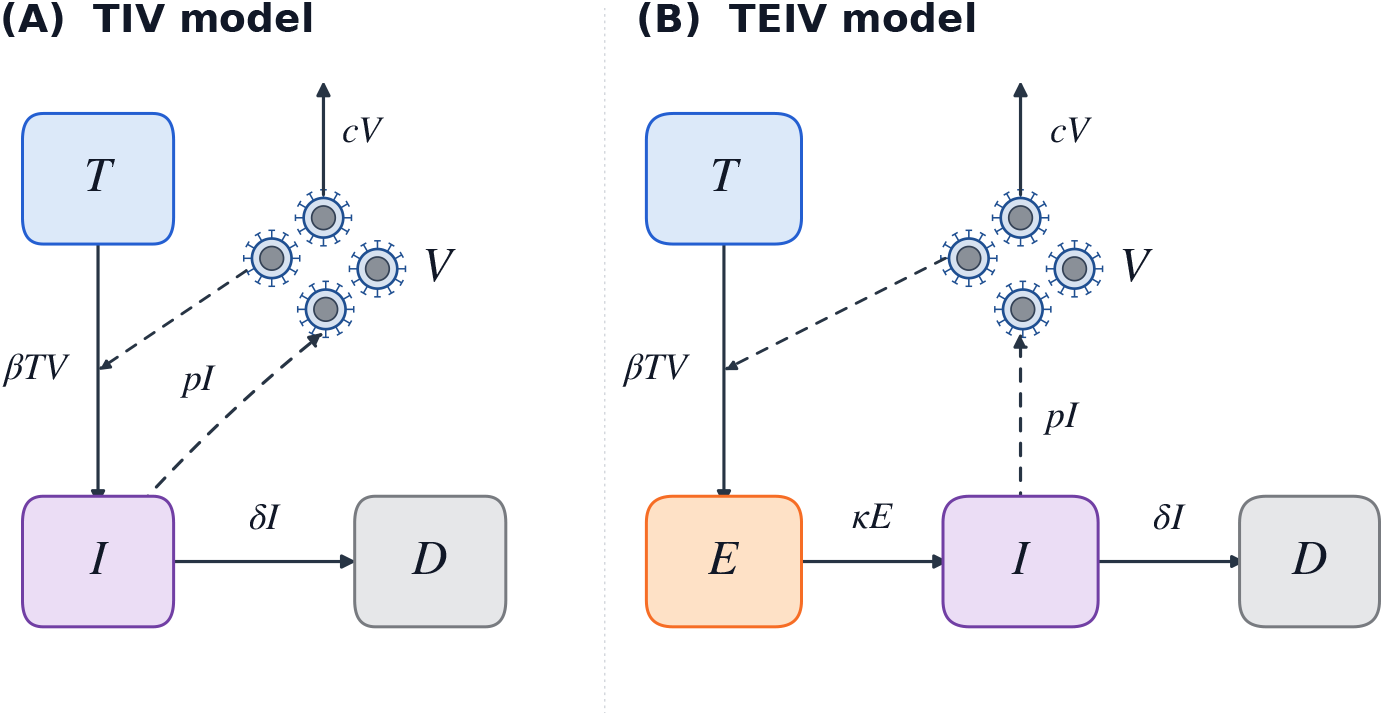
Flow diagrams of the TIV (A) and TEIV (B) models defined by systems (2) and (3), respectively. *T* denotes susceptible target cells, *E* non-productively infected cells, *I* productively infected cells, *D* dead infected cells, and *V* viral RNA concentration.

**Table 1:**
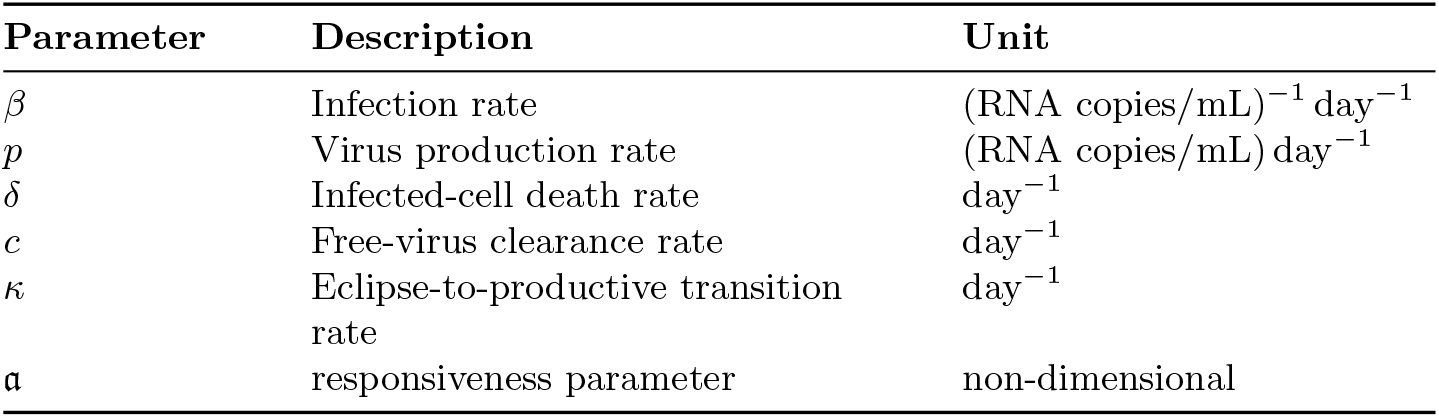
Parameters of the models. The first four appears in all models, *κ* is only in TEIV, and *a* is only in fr-TIV.

#### 2.3.2 TEIV model

The TEIV model extends TIV by introducing a non-productively infected eclipse compartment (*E*) between target-cell infection and productive infection [10]. Newly infected cells therefore enter *E* before transitioning to the virus-producing compartment *I*.

The TEIV equations are obtained by extending the TIV system (2):

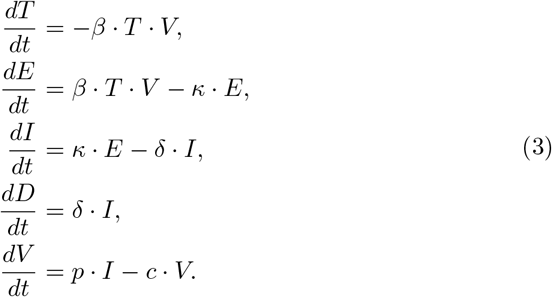

In the TEIV model, target cells enter the eclipse phase at rate *βTV*, and cells remain in compartment *E* for an average of *κ^−^*^1^ days before becoming productively infected. Productively infected cells release virus at rate *p*, die at rate *δ*, and free virus is cleared at rate *c*.

The interactions in the TEIV model are illustrated in panel B in Figure 1.

The model parameters are summarized in Table 1.

#### 2.3.3 Fractional-order TIV model (fr-TIV)

The fractional-order TIV model (fr-TIV) extends the classical TIV formulation using Caputo–Fabrizio fractional operators of order *a ∈* (0, 1) [16]. Unlike the TEIV model, which introduces an additional biological compartment, the fractional-order formulation instead modifies the temporal evolution of the existing compartments through a fractional derivative. Fractional-order models have been increasingly applied to biological and epidemic systems because they provide a flexible framework for describing processes with memory and hereditary effects that are not captured by classical integer-order models [16]. In such models, the current dynamics depend not only on the present state but also on the system’s past trajectory, allowing history-dependent processes to be represented naturally.

Because time is measured in days, we use dimensionally scaled Caputo– Fabrizio operators with the fixed reference timescale *t*_ref_ = 1 day [20]. This reference timescale is not fitted; its numerical value is 1 in the implemented time unit.

For any real, absolutely continuous function *f,* the Caputo–Fabrizio [21] fractional derivative of order 0 *< a <* 1, is defined by

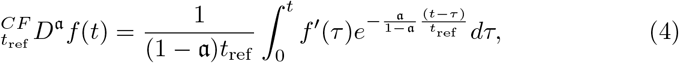

while the corresponding fractional integral is given by

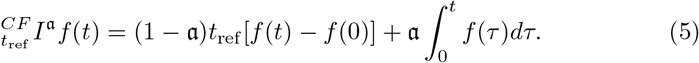

Let *T, I, D* denote the number of susceptible target cells, infected cells and dead cells, respectively, and let *V* denote the virus concentration. As introduced in [22], we formulate the Caputo–Fabrizio fractional-order extension of the TIV model as follows:

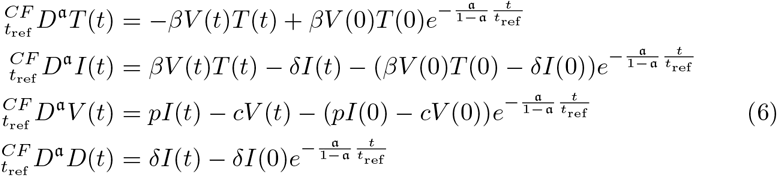

with fractional derivative order *a ∈* (0, 1) and parameters *β, δ, p, c >* 0 representing the cell infection rate, cell death rate, cellular virus production rate, and viral clearance rate, respectively. For the initial conditions, we have

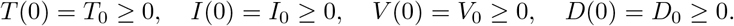

The exponentially decaying correction terms in (6) are included to satisfy the Caputo–Fabrizio compatibility condition at the initial time: since *^CF^ D^a^f* (*t*) *→* 0 as *t →* 0+ for admissible functions, the right-hand side of each fractional equation must vanish at *t* = 0.

As detailed in [22]; for sufficiently regular solutions, system (6) is equivalent to the following implicit first-order ODE system:

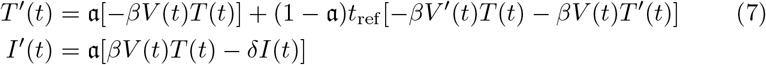

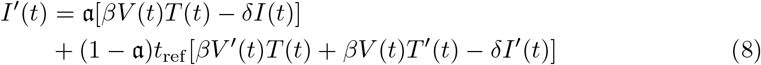

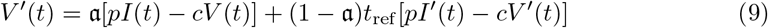

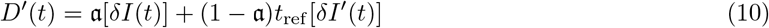

with initial conditions

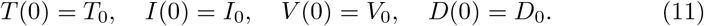

This system is linear in the derivatives. Whenever the regularity determinant is nonzero, it can be solved algebraically for *T ^′^*, *I^′^*, *V ^′^*, and *D^′^*; we verified this symbolic elimination using SymPy and used the resulting explicit vector field for numerical integration.

From the ODE form, we can interpret *a* as a responsiveness parameter, see [22]. For numerical model selection we restricted this parameter to *a <* 1, rather than allowing the endpoint *a* = 1, because this endpoint coincides with the classical TIV model.

### 2.4 Initial conditions and parameter intervals

Table 2 summarizes the initial values and parameter intervals we used for optimization.

**Table 2:** Initial conditions and implemented parameter intervals used for model fitting.

| Parameter | Implemented interval | Source |
| --- | --- | --- |
| $T_0$ | dimensionless, fixed to 1.0 | Target-cell-related state variables were normalized to the initial target-cell level for fitting. Thus $T_0 = 1$ is a scale convention, not an independently estimated biological cell count. |
| $V_0$ | 1.0 copies/mL, fixed | Fixed to a value of 1.0 for numerical simplicity. |
| $I_0$ | 0, normalized units | Assumed to be zero at the start of infection. |
| $E_0$ | 0, normalized units | Assumed to be zero at the start of TEIV simulations. |
| $D_0$ | 0, normalized units | No infected-cell deaths have accumulated at the start of the simulation. |
| $\beta$ | $[10^{-9}, 10^{-2}]$ | Broad optimization interval for the infection-rate parameter. The interval given in [10] could not be adopted due to the assay-specific nature of $\beta$ |
| $R_0$ | $[1.01, 100]$ | Dimensionless reproduction-number parameter optimized instead of fitting the production rate $p$ independently. |
| $p$ | Derived, not independently fitted | For each candidate parameter vector, the production rate was calculated from $p = R_0 c \delta / \beta$, using the corresponding model's $\beta$, $c$, and $\delta$. |
| $c$ | 2.4 day<sup>-1</sup>, fixed | Fixed to the value used by [10]. |
| $\kappa$ | $[0.5, 4]$ day<sup>-1</sup> | [18] and [23]. |
| $\delta$ | $[0.61, 4.0]$ day<sup>-1</sup> | Union of the reported 95% confidence intervals in [10], extended to improve fit. |
| $\mathfrak{a}$ | $[0.2, 0.999]$ | The upper bound excludes the endpoint $\mathfrak{a} = 1$, where the model coincides with classical TIV. |

Although the TIV model structure and the type of Ct observations in the present paper are similar to those of [10], we highlight that we did not directly adopt their fitted production (*p*) and infection-rate (*β*) parameters. In particular, the fitted *β* is an effective infection-rate coefficient on the swab-measurement scale: it absorbs unmodelled factors relating local airway viral abundance to the assayed RT-qPCR sample, such as sampling efficiency, dilution, PCR calibration, and laboratory processing. The dataset used in [10] and the NBA dataset analyzed here were obtained using different sampling and assay protocols. Consequently, the search bounds for *β* were re-estimated, and they should be interpreted as a functional optimization range rather than as direct microscopic virus–cell infectivity constants. The bounds are biologically motivated only at the order-of-magnitude/effective-model level, and are deliberately broad because *β* depends on the measurement scale.

At the start of each simulation, the system contains only target cells and an initial viral inoculum. Thus, *I*_0_ = *D*_0_ = 0 in all models and *E*_0_ = 0 in TEIV, while *T*_0_ and *V*_0_ are fixed as specified in Table 2.

### 2.5 Parameter estimation and optimization

Each retained infection wave was fitted separately with the TIV, TEIV, and fr-TIV models. Parameter estimation minimized the profiled negative log-likelihood under the right-censored Gaussian observation model defined below [24, 25].

The continuous model parameters were optimized numerically, whereas the integer-valued time offset *τ* was handled by exhaustive enumeration over its biologically admissible range. Specifically, observation *i* was compared with the model prediction at shifted model time *t_i_* = *x_i_* + *τ*. Negative offsets were allowed only when they shifted leading Ct= 40 measurements before the model’s infection start; no Ct value below 40 was allowed to precede model time zero, i.e., infection start. Positive offsets were allowed up to 28 days, a permissive upper bound for acute waves relative to the typical detectability window of SARS-CoV-2 RNA, which is reported to be cca. 21 days [26]. For shifted sampling times preceding the model start, the viral load was taken to be zero; under the Ct observation link, this was represented by the limit *µ_i_ →* +*∞*, for which the probability of a recorded Ct value of 40 is one. For the information-criterion penalty, *τ* was counted as one fitted parameter.

The fitted parameters were (*β, R*_0_*, δ, τ*) for TIV, (*β, R*_0_*, δ, κ, τ*) for TEIV, and (*β, R*_0_*, δ, a, τ*) for fr-TIV. The viral-clearance rate and initial conditions were fixed, and the viral production rate *p* was derived from the fitted parameters; these quantities were therefore not counted as fitted parameters.

The global search was performed using Nevergrad [27]. We used default parameter values and Latin hypercube samples to initialize the search. For every continuous parameter vector proposed during optimization, the model trajectory was simulated once, and the residual standard deviation *σ* was profiled separately at every admissible integer offset to obtain an offset-specific negative log-likelihood. The smallest of these offset-specific objective values was returned to the optimizer, while the values for all offsets were retained. Thus, no single offset was fixed during the global search.

After the Nevergrad phase, the best previously evaluated parameter vectors were selected as starting points for local optimization. These starting points were refined using the Nelder–Mead algorithm implemented in SciPy [28]. During local refinement, every new continuous parameter vector was again evaluated over the complete admissible offset range; consequently, the best-fitting offset could change during a Nelder–Mead run.

The initial optimization stage used 20 Latin hypercube points, a Nevergrad budget of 300 evaluations, and up to six local refinements. We then assessed whether the best solution was numerically confirmed. Let *Q*_min_ denote the lowest profiled negative log-likelihood found. A solution was considered confirmed if at least one successfully converged local refinement had an objective value within 10*^−^*^3^ max(*|Q*_min_*|,* 1) of *Q*_min_, or if at least two separate local refinements produced mutually consistent near-best objective values under the same scaled tolerance. If neither condition was met, the search was automatically repeated with an independent deterministic seed, using 48 Latin hypercube points, a Nevergrad budget of 1000 evaluations, and up to 12 local refinements. The lowest objective value found across all completed stages was always retained. If the result remained numerically unconfirmed, the three most promising offsets identified during the preceding searches were fixed in turn, and the continuous parameters were refined separately at each offset. If a fit still could not be numerically confirmed after this final step, the best solution found would have been kept in the analysis and marked as numerically unconfirmed. In the present analysis, however, all model fits were numerically confirmed.

All random components used reproducible deterministic seeds. Optimization diagnostics, including the selected offset, convergence information, and the escalation stage reached, were stored for every model fit. The code is available at [29].

### 2.6 Likelihood and model-selection criteria

#### 2.6.1 Censored-Gaussian likelihood

An infection wave is represented by *n* observations (*x_i_, c_i_*), where *x_i_* is the sampling time and *c_i_* is the recorded Ct value. Let *µ_i_* denote the corresponding untruncated Ct-scale mean predicted by the model, as defined above. We assumed an unobserved latent measurement

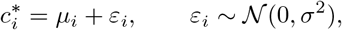

with independent measurement errors [24, 25]. The recorded value *c_i_* was related to the latent measurement by

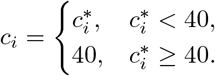

Thus, a recorded value below 40 was treated as an uncensored observation, whereas a recorded value of 40 indicated only that the unobserved latent Ct measurement was at least 40. The residual standard deviation *σ* was estimated separately for each model fitted to each infection wave.

For model *M*, the log-likelihood for one infection wave was

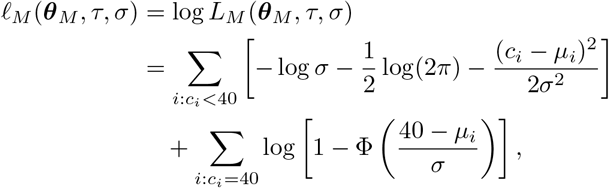

where Φ denotes the cumulative distribution function of the standard normal distribution.

Because AICc was calculated from the fitted censored-Gaussian likelihood, we assessed the adequacy of the associated error assumptions using diagnostics of residual normality, time-dependent bias, variance stability, and serial dependence. The residual behavior was broadly compatible with the normal approximation and no severe departures were identified, supporting the likelihood as a reasonable working approximation (see Supplementary Notes).

#### 2.6.2 Information criteria

The information criteria were calculated from the maximized log-likelihood under the censored Gaussian observation model defined above.

Let 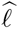 denote the maximized log-likelihood for a candidate model fitted to one infection wave. AIC [30] was calculated as

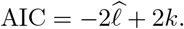

Here, *k* denotes the total number of estimated parameters, including the fitted time offset *τ* and the measurement-error standard deviation *σ*. Thus, *k* = 5 for TIV and *k* = 6 for TEIV and fr-TIV. The sample size *n* was the number of Ct observations included in the fitted wave, including observations recorded at *c_i_*= 40.

Lower AIC values indicate stronger support among the candidate models.

Because the number of observations within an infection wave can be small relative to the number of fitted parameters, our primary model-selection criterion is the small-sample corrected Akaike information criterion (AICc) [31],

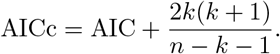

AICc is defined only when *n > k* + 1. Lower AICc values indicate stronger relative support among the candidate models. Uncorrected AIC is retained as a sensitivity analysis.

The Bayesian information criterion (BIC) [32] was used as a second sensitivity analysis:

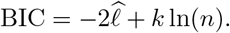

None of these criteria measures absolute goodness of fit; they are intended for relative comparison among models fitted to the same infection wave.

For the categorical summaries reported below, models with information-criterion values within 2 units of the within-wave minimum (ΔIC *≤* 2) were classified as effectively tied [10, 33]; otherwise, the minimum-IC model was classified as the clear winner.

## 3 Results

### 3.1 Illustrative fitted trajectories

We first show an example in which all three candidate models closely reproduce the observed Ct trajectory. In the wave shown in Figure 2, TEIV achieved the lowest negative log-likelihood (NLL), whereas TIV was the clear AICc winner. We next examine the trade-off between a lower unpenalized NLL and the AICc penalty for additional model complexity.

**Figure 2:**
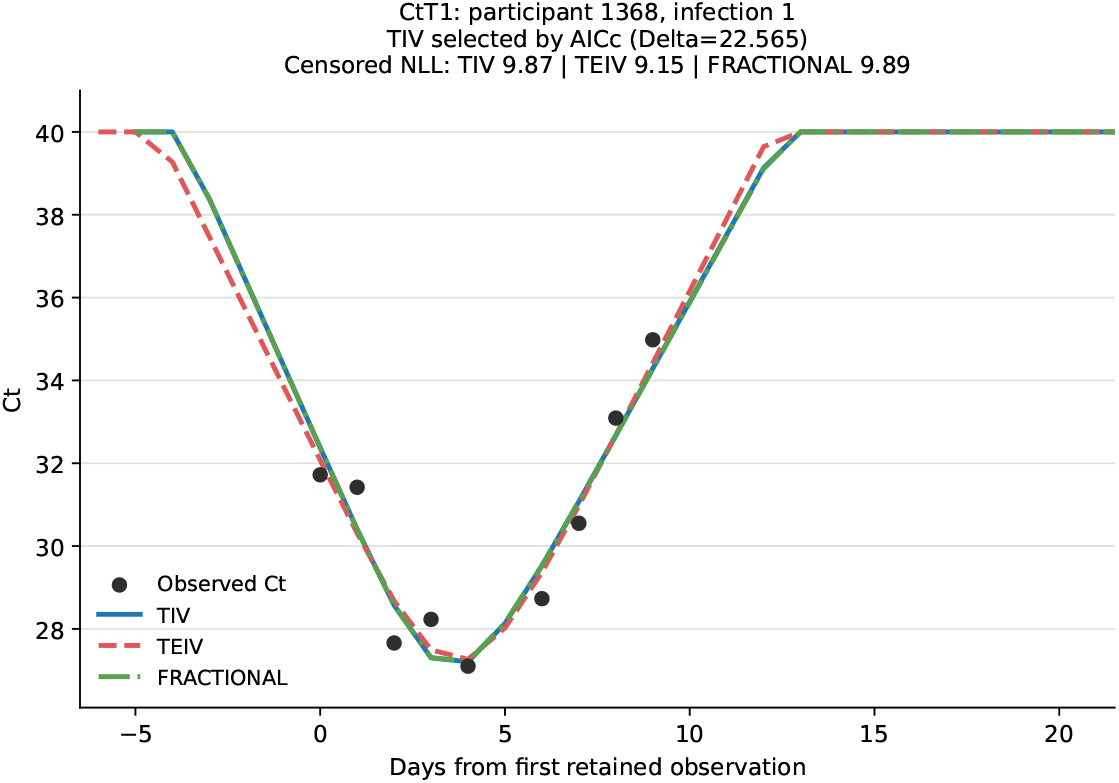
Illustrative fitted trajectory for participant 1368, infection 1. Points denote the observed Ct measurements and lines show the fitted trajectories. TEIV achieved the lowest NLL, while TIV was clearly selected by AICc. The wave was selected as the case with the smallest TEIV NLL among waves satisfying these two conditions.

Figure 3 presents two infection waves chosen to illustrate contrasting relationships between goodness of fit and AICc-based model selection. In panel A, the reduction in the negative log-likelihood is sufficiently large to support the additional flexibility of the fractional-order model. In panel B, the fractional-order model also attains a lower negative log-likelihood than TIV, but this improvement is not sufficient to overcome the small-sample complexity penalty, resulting in an effective AICc tie between the two models.

**Figure 3:**
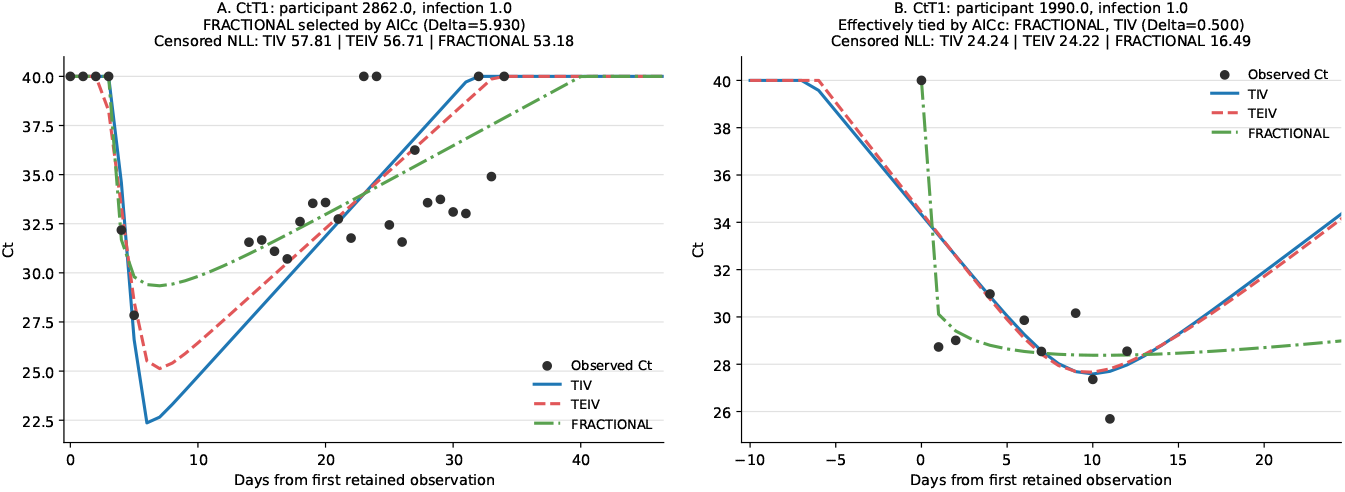
Illustrative fits of the TIV, TEIV, and fr-TIV models to two CtT1 infection waves. Black points denote the observed Ct measurements and lines show the fitted trajectories. In panel A, the substantial improvement in likelihood-based fit provided by the fr-TIV model is sufficient to justify its additional parameter, and the model is clearly selected by AICc. In panel B, the fr-TIV model achieves a lower negative log-likelihood than TIV, but the improvement is insufficient to offset the AICc complexity penalty; TIV and fr-TIV are there-fore effectively tied. The examples were chosen to illustrate these contrasting model-selection outcomes.

Together, these examples show why goodness of fit and model selection must be considered separately. AICc can support an additional parameter when it produces a sufficiently large improvement in fit, as in panel A, while a smaller improvement may not justify the added complexity, as in panel B.

The frequency of these contrasting model-selection outcomes across all retained infection waves is examined in the following aggregate analysis.

### 3.2 Aggregate model performance

This section presents aggregate model rankings based on both the unpenalized likelihood and information criteria that account for model complexity.

The unpenalized censored-likelihood ranking is shown in Figure 4. In this ranking, models with a fitted likelihood of at least 99% of the within-wave maximum were considered effectively tied whenever more than one model met this threshold; otherwise, the model with the highest fitted likelihood was the clear winner. TIV was the clear best-fitting model in 29 of 868 evaluable CtT1 trajectories (3.3%), TEIV in 295 (34.0%), and fr-TIV in 262 (30.2%); the remaining 282 trajectories (32.5%) were effectively tied. Thus, before accounting for model complexity, the additional flexibility of TEIV and fr-TIV often yielded better within-sample fit than TIV.

**Figure 4:**
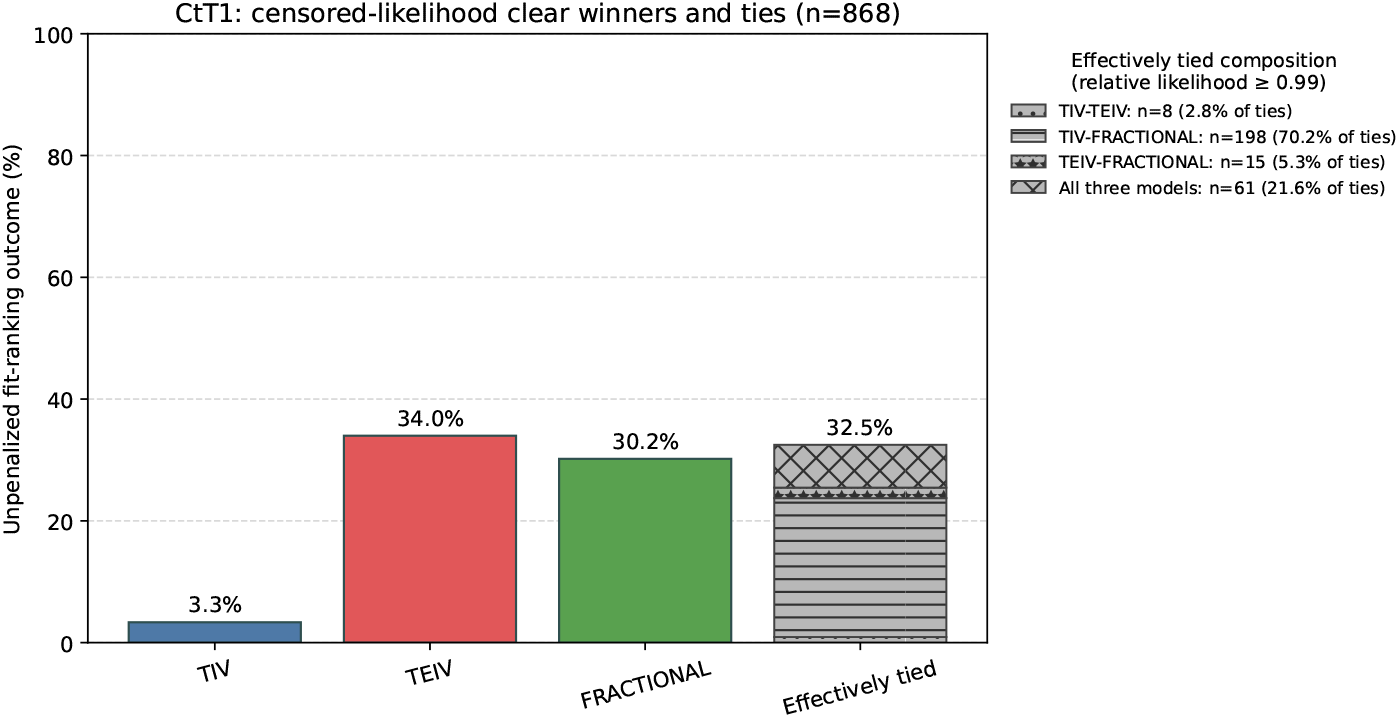
Percentages of clear winners and effective ties based on the unpenalized likelihood under the censored Gaussian observation model. The stacked tie bar shows which models formed the tied sets.

The AICc ranking shown in Figure 5 reflects the combined effects of the maximized log-likelihood, the number of fitted parameters, and the small sample size of each trajectory. It differs substantially from the unpenalized-likelihood ranking. Under AICc, TIV was the clear winner in 849 of 868 Ct trajectories (97.8%), fr-TIV in 4 (0.5%), and TEIV in none; the remaining 15 trajectories (1.7%) were effectively tied. Thus, in most trajectories, the likelihood improvement achieved by the more complex models was insufficient to offset the AICc penalty associated with their additional fitted parameter.

**Figure 5:**
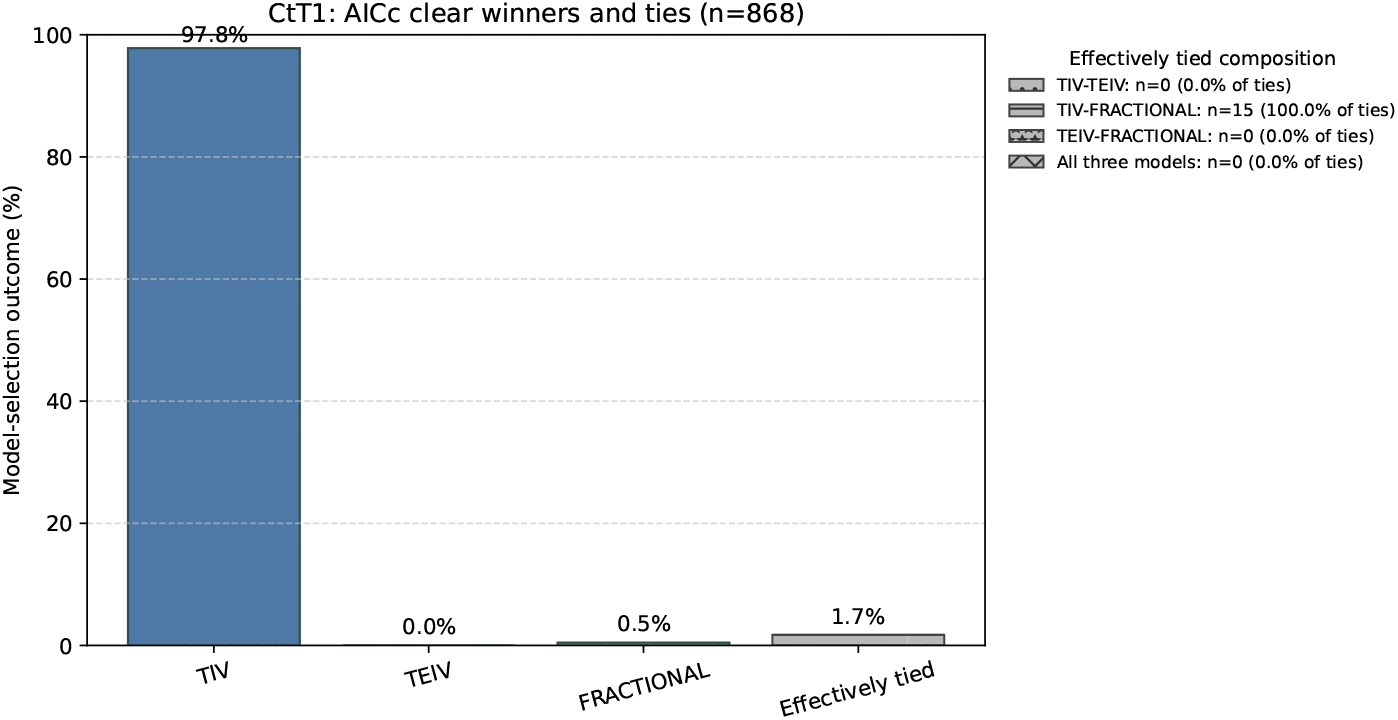
Percentages of clear winners and effective ties according to AICc.

We repeated the aggregate ranking using AIC and BIC as sensitivity analyses, since neither criterion includes the small-sample correction applied in AICc. The corresponding results are shown in Figures 6.

**Figure 6:**
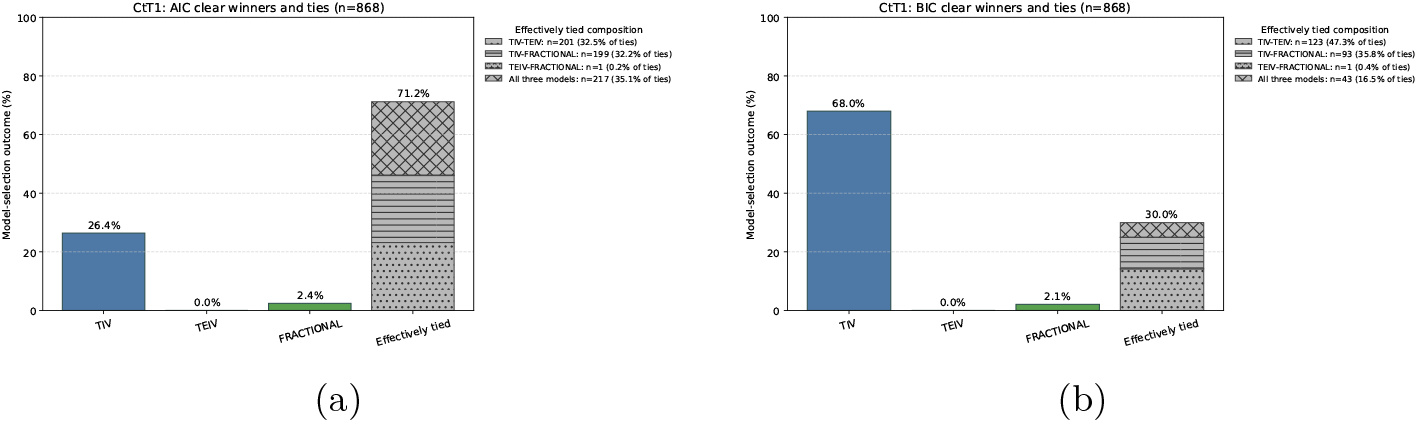
Percentages of clear winners and effective ties according to AIC (left) and BIC (right).

The distributions of the gaps between the best- and second-ranked models shown in Figure 7 quantify the degree of model separation underlying the categorical clear-winner and tie summaries.

**Figure 7:**
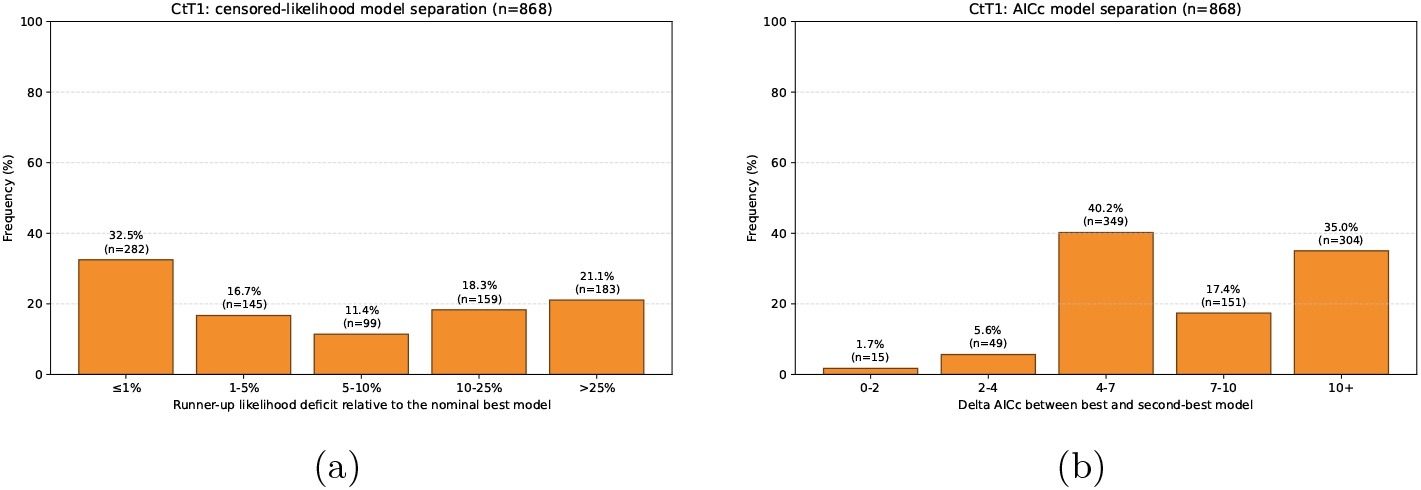
Distribution of within-wave separation between the nominal best and runner-up models according to the unpenalized censored-Gaussian likelihood (left) and AICc (right). The likelihood deficit was calculated as 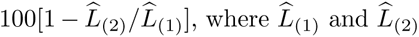 are the largest and second-largest fitted likelihoods; the AICc gap was calculated as AICc_(2)_ *−* AICc_(1)_. Bars show percentages of infection waves. Likelihood deficits of at most 1% and AICc gaps of at most 2 were classified as effective ties.

### 3.3 Sensitivity to the Ct-to-viral-load conversion

Ct-to-viral-load standard curves are established through dedicated wet-lab calibration experiments (e.g., serial dilutions of a synthetic RNA transcript of known concentration) and are therefore assay- and target-specific. Consequently, they are not universally available. Even when two assays use the same PCR platform and nominal target, the slope and intercept of the Ct–concentration relationship may differ because they depend on the reference material, amplification efficiency, sample matrix, extraction and volume conventions, and fluorescence-thresholding procedure.

A curve calibrated for one assay or target cannot generally be assumed to hold for another. It is therefore useful to know whether downstream modeling conclusions are sensitive to the specific choice of conversion.

As noted before, the CtT1 values in NBA data [12] were generated with the Roche cobas Target 1 assay whereas the Zhang et al. [17] conversion was calibrated for ORF1ab Ct measurements. The Zhang conversion formula is given by

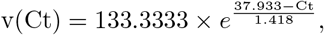

while the Kissler conversion formula is defined in equation (1). Both conversions belong to the same log-linear family, log_10_ *v* = *a − b* Ct: we used the Kissler calibration (*a, b*) = (15.470, 0.371) as the primary observation link and the Zhang calibration (*a, b*) = (13.743, 0.306) as a published alternative for assessing the sensitivity of the model-selection results to changes in *a* and *b*. This corresponds to a moderate but non-negligible 18% change in slope.

To assess whether the choice between the Kissler [13] and Zhang [17] Ct-to-viral-load conversions materially affects fit quality, we compared the results obtained under each conversion. As a visual example, we show one wave and fit for each conversion formula in Figure 8.

**Figure 8:**
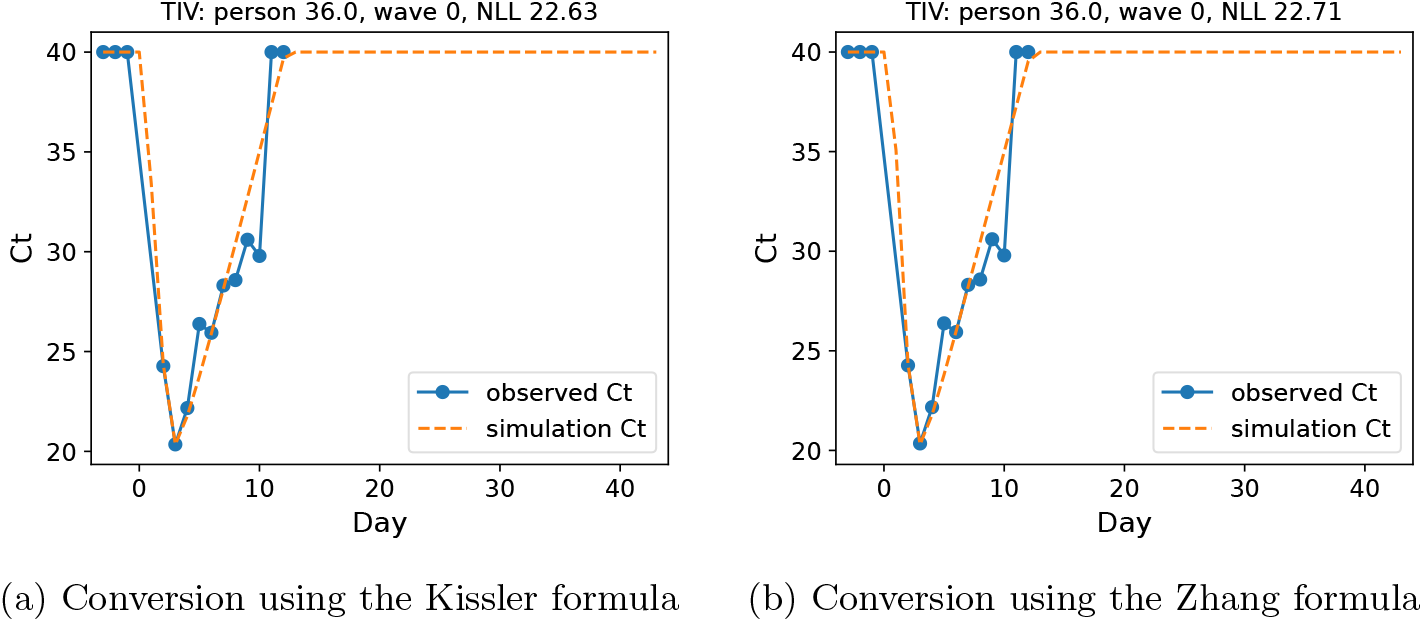
Individual waves shown with Kissler and Zhang conversion formula, respectively. The negative log-likelihoods are shown in the graphs.

Under both Ct-to-viral-load conversions, all three models – including the simplest TIV model – reproduced the Ct time series similarly well. Thus, the change introduced by the conversion was largely absorbed by re-estimating the parameters, without requiring a more complex dynamical structure. These conversion-associated parameter changes are illustrated for *β*_TIV_ in Figure 9 by comparing paired estimates obtained for the same waves under the Kissler and Zhang conversions. The TEIV and fractional models showed similar paired patterns.

**Figure 9:**
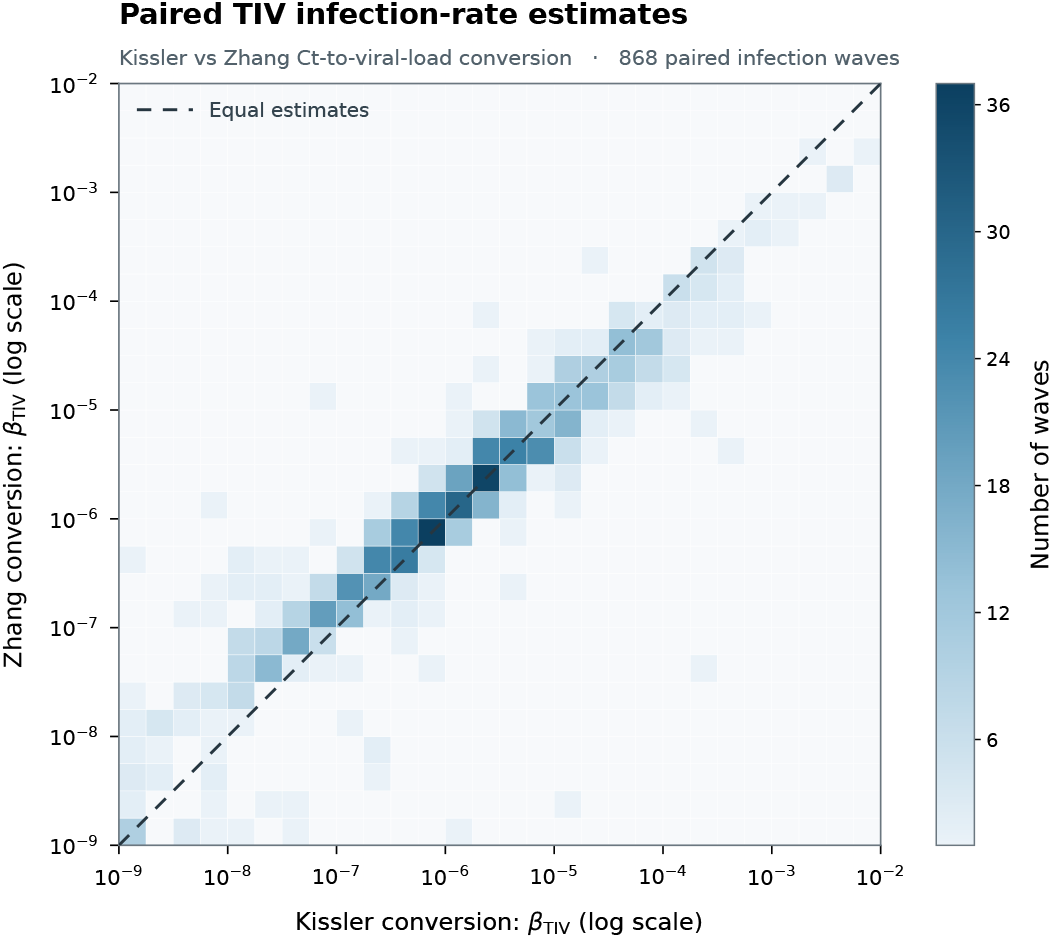
Paired TIV infection-rate parameter estimates under the Kissler [13] and Zhang [17] Ct-to-viral-load conversions.

This supports our main finding that the TIV model is already sufficiently flexible on its own to describe the observed trajectories, and that its relative performance is not a consequence of a particular or perfect Ct-to-viral-load conversion.

We note that the several-orders-of-magnitude dispersion of *β* shown in Figure 9 is likely to reflect a combination of observation-scale variability, genuine between-wave heterogeneity, parameter compensation, and limited practical identifiability.

Model-selection outcomes under the two conversions are shown in Figure 10. Exact agreement was 97.8% under AICc and 69.0% under the unpenalized censored likelihood; among the 269 likelihood-based disagreements, 177 involved a tie under one conversion, whereas 92 of 868 waves (10.6%) yielded different clear winners. Overall, the choice of conversion has little effect on AICc-based model selection, although the unpenalized likelihood ranking is more sensitive.

**Figure 10:**
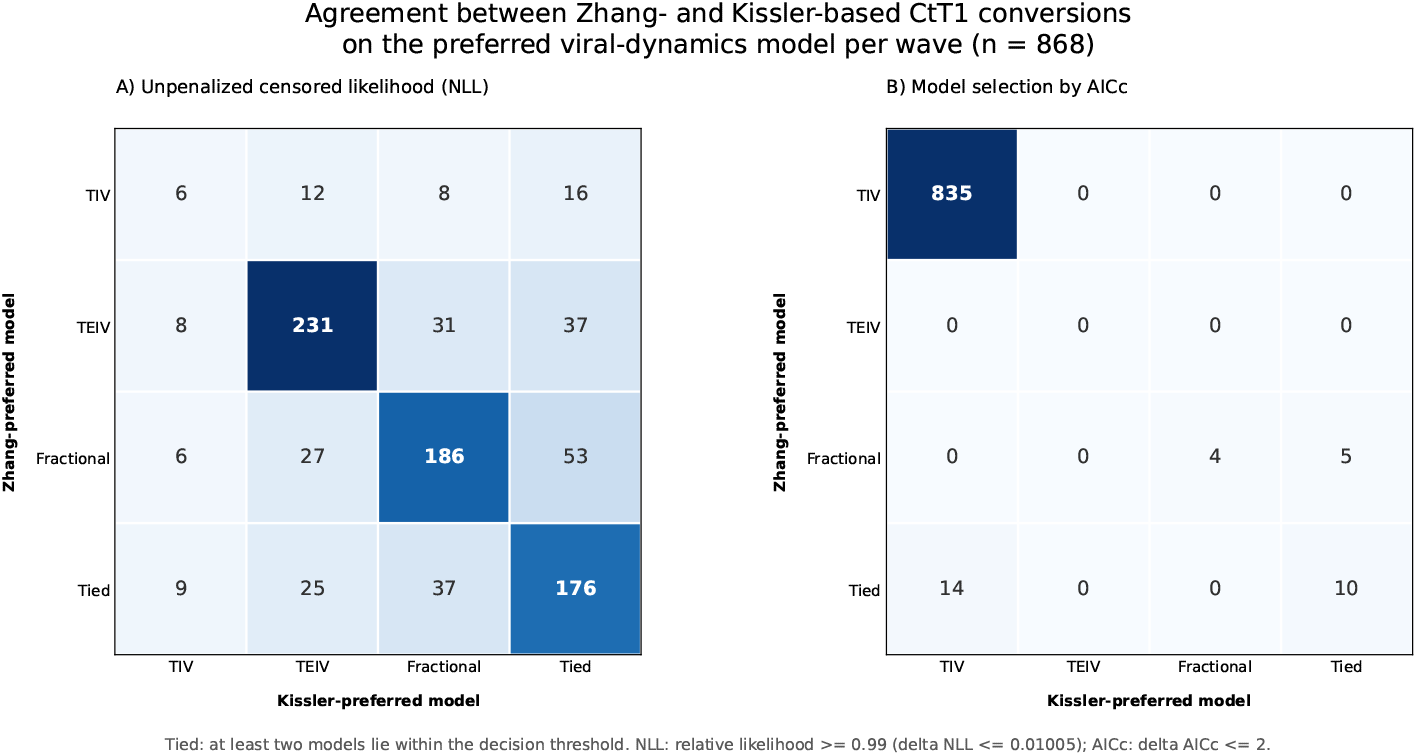
Agreement in model-selection outcomes between the Zhang and Kissler Ct-to-viral-load conversions under the unpenalized censored-Gaussian likelihood (left) and AICc (right).

## 4 Discussion

Several mathematical formulations can be used to describe within-host SARS-CoV-2 dynamics, each emphasizing different biological mechanisms. This study examined how to select among competing models by considering both goodness of fit and model complexity. We analyzed longitudinal viral-testing data from NBA participants and reconstructed temporally localized groups of observations as infection waves.

Three models were fitted to every retained wave: TIV, TEIV, and fr-TIV. Each extends the target-cell-limited framework by adding an eclipse phase or fractional-order dynamics. Increasing complexity allows more detailed trajectories to be represented. Unpenalized fit was evaluated using the negative log-likelihood, while model selection was based on AICc, AIC, and BIC computed from the maximized log-likelihood.

The negative log-likelihood may decrease as the number of parameters increases, because additional flexibility can permit closer agreement with the data. AICc, AIC, and BIC, however, penalize this complexity. Model selection should therefore consider not only in-sample accuracy but also parsimony and the ability to generalize.

Across the evaluable CtT1 trajectories, AICc predominantly selected TIV. The more flexible models were able to improve likelihood-based fit, but these improvements were generally insufficient to justify their additional parameter. The resulting preference for TIV should therefore be interpreted as support for a parsimonious effective representation of the information contained in the observed Ct trajectories.

The results should be interpreted in the context of the short duration and relatively sparse sampling of the analyzed infection waves. The information content of the available Ct trajectories did not justify the separate parameterization of additional mechanisms beyond those represented by the TIV model. Among the candidate models, TIV was therefore the best-supported effective description of the observed dynamics. Importantly, the limited support for TEIV should not be interpreted as evidence that an eclipse phase is biologically absent. Rather, under the present sampling design, the Ct-only trajectories provided insufficient information to support the eclipse phase as a separately estimable component of the model.

The contrast between rankings based on negative log-likelihood and AICc is central to the interpretation of our results. Although the additional flexibility of the TEIV and fractional-order models often improved likelihood-based fit, these improvements were generally too small to justify the additional parameter. TIV was therefore the best-supported description for the short, sparsely sampled, Ct-only trajectories analyzed here. This is a data- and purpose-specific model-selection result, rather than evidence that eclipse-phase or memory-like dynamics are biologically absent.

More generally, our results emphasize that model complexity should be matched to the mechanistic resolution supported by the available observations. Distinguishing eclipse-phase or memory-like dynamics is likely to require dense sampling or complementary biological measurements beyond Ct trajectories alone.

Concerning limitations, we mention that target-cell-limited models can reproduce viral decline by depleting the susceptible-cell compartment [10, 34]. Such depletion should not be interpreted literally as extensive epithelial-cell loss in vivo. Thus, the AICc preference for TIV in this analysis supports a parsimonious description of the observed Ct trajectories, rather than proving that target-cell depletion is the sole biological driver of viral clearance.

As a direction for future work, we highlight the Ct T2 data in the dataset. For now, this was not taken into consideration, as target 2 works with a different gene than target 1, and no standard curve is available for conversion.

## Acknowledgements

GR, NJ, DS, DRC and BD were supported by 2019-2.1.11-TÉT-2020-00231. DS and DRC gratefully acknowledge the financial support of the Ministry of Science, Technological Development and Innovation of the Republic of Serbia (Grants No. 451-03-33/2026-03/200125 and 451-03-34/2026-03/200125). GR, FAB and NJ were supported by 2024-1.2.3-HU-RIZONT-2024-00034. NJ was supported by Project No. 152347, funded by the Ministry of Science and Technology of Hungary through the National Research, Development and Innovation Fund under the STARTING 25 funding scheme.

