## Supplementary material for "Fitting and Selection of Within-Host Models for SARS-CoV-2 Infection"

N. Juhász, D. Selesi, D. Nagy, B. Das, F. A. Bartha,
D. Rajter-Ciric and G. Röst

### 1 Residual diagnostics

The AICc comparison was based on the maximized log-likelihood of a censored-Gaussian observation model. We therefore assessed whether, within each separately fitted model–wave combination, the latent Ct-scale measurement errors could reasonably be approximated as

$$\epsilon_i \overset{\text{iid}}{\sim} N(0, \sigma^2),$$

where $\sigma$ was estimated separately for each fit. The aim was not to prove exact Gaussian distributions, but to identify possible departures sufficiently large or systematic to make the Gaussian observation model questionable.

Formal tests of exact normality were not used as the primary assessment. This is in part because their conventional calibration does not directly accommodate censoring. We therefore used graphical diagnostics. Together, for $Z_i = \epsilon_i/\sigma$, these graphical diagnostics addressed four aspects of the working error model:

1. pooled agreement with the standard-normal reference;
2. centering around zero over the infection wave, i.e. $E(Z \mid \text{wave phase}) \approx 0.$;
3. stability of residual spread, assessed by $\sqrt{E(Z^2 \mid \text{wave phase})} \approx 1$; and
4. a necessary implication of temporal independence, assessed through $E(Z_t Z_{t+d}) \approx 0$ at short lags.

Because individual waves contained too few observations for reliable fit-specific residual diagnostics, the diagnostics were evaluated in aggregate for each candidate model after standardization by the fit-specific residual scale.

The pooled diagnostics were intended to identify systematic departures shared across waves rather than to establish the error distribution of every individual fit.

### 1.1 Normal Q–Q assessment

Normality of the Ct-scale measurement errors was assessed separately for each candidate model using censoring-aware randomized quantile residuals [1] pooled across the fitted infection waves. For observations below Ct 40, the ordinary standardized residual was used, whereas observations recorded at Ct 40 were randomized over the conditional upper tail implied by the fitted censored-Gaussian model. The residual quantiles follow the normal reference line with modest deviations in the extreme tails and no gross departure; the shaded region represents variation due to the randomization of censored observations and it is not a sampling-based confidence interval.

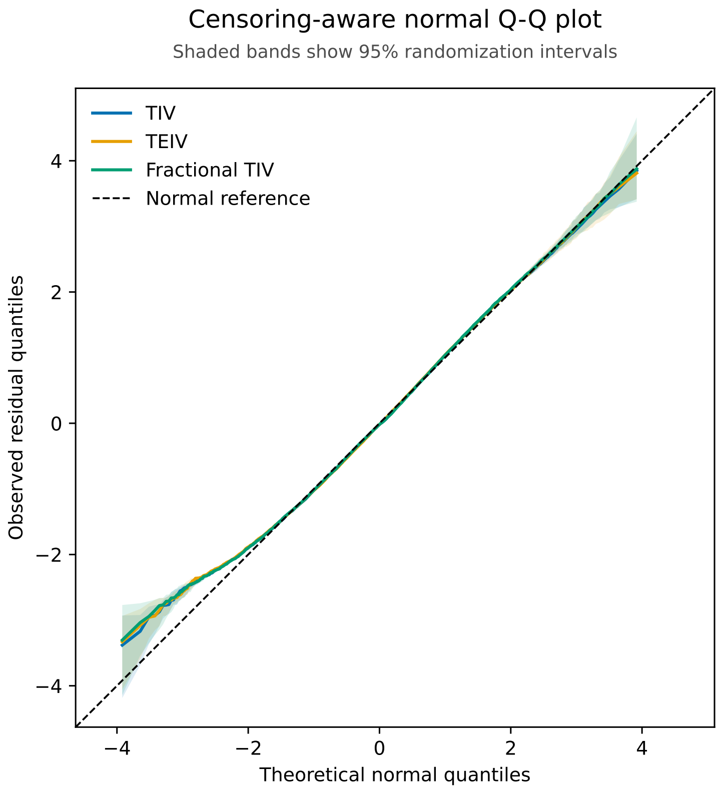


Figure 1: Normal Q–Q assessment of the censoring-aware residuals for the TIV, TEIV, and fractional-order TIV models. The diagonal denotes the standard-normal reference, and the shaded region shows the central 95% of the results obtained by re-randomizing the censored observations.

### 1.2 Mean residual over time

In order to examine time-dependent mean bias, measurements were grouped into five periods according to the number of days elapsed since the first observation in each wave. For uncensored observations, the ordinary standardized residual was used; for censored observations, it was replaced by its conditional expectation under the fitted model, i.e.

$$\widetilde{Z}_i = \begin{cases} \dfrac{C_i - \widehat{\mu}_i}{\widehat{\sigma}_i}, & C_i < 40, \\ E(Z_i \mid Z_i \geq a_i) = \dfrac{\phi(a_i)}{1 - \Phi(a_i)}, & C_i = 40, \end{cases} \qquad a_i = \frac{40 - \widehat{\mu}_i}{\widehat{\sigma}_i},$$

where $\phi$ and $\Phi$ denote the standard-normal density and distribution function, respectively. For each period $b$, the plotted value was $\overline{Z}_b = \frac{1}{n_b}\sum_{i \in b} \widetilde{Z}_i$. The period-specific means therefore provide a coarse, pooled assessment of $E(Z \mid \text{wave phase}) \approx 0$, rather than a pointwise or fit-specific test. The vertical bars show central 95% intervals from 2,000 participant-level resamples of the already fitted residuals; the models were not refitted. Across all models, no large systematic early-to-late mean bias was apparent. The result is broadly consistent with the working assumption of no systematic time-dependent mean bias.

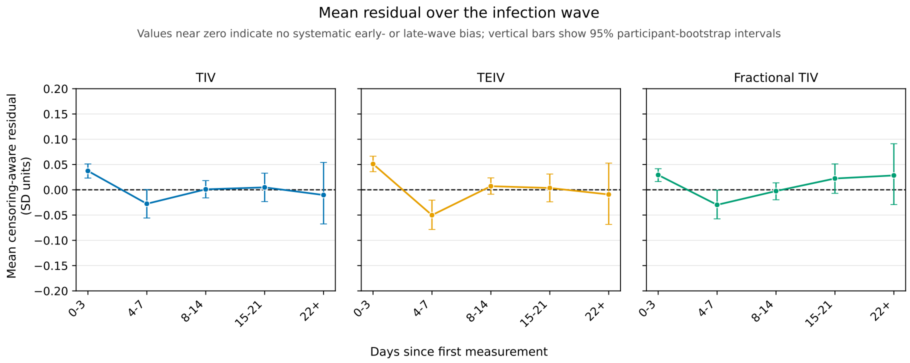


Figure 2: Mean standardized residuals over the fitted infection waves. Points show the period-specific means, and vertical bars show central 95% participant-bootstrap intervals.

### 1.3 Residual spread over time

Measurements were grouped into seven periods to examine whether residual spread changed over time. For each observation, we defined

$$m_i = \begin{cases} \left(\dfrac{C_i - \widehat{\mu}_i}{\widehat{\sigma}_i}\right)^2, & C_i < 40, \\ E(Z_i^2 \mid Z_i \geq a_i) = 1 + a_i \dfrac{\phi(a_i)}{1 - \Phi(a_i)}, & C_i = 40, \end{cases}$$

where $a_i = (40 - \widehat{\mu}_i)/\widehat{\sigma}_i$. For period $b$, the plotted residual spread was

$$R_b = \left( \frac{1}{n_b} \sum_{i \in b} m_i \right)^{1/2}.$$

Under the working observation model, the reference value is $R_b = 1$. Because the period-specific residual means were close to zero, this quantity reflects residual spread within each period.

The residual spread remained close to 1 through day 19. From day 20 onward, the residual-spread estimates showed a clear but moderate upward pattern; they increased to approximately 1.15–1.17 during days 20–23 and to 1.20–1.21 from day 24 onward across the three models. This is consistent with somewhat greater late-wave residual spread than represented by the fitted constant-$\sigma$ model. However, because the final period contained many censored measurements and the composition of the late-period group is also determined by the Ct-based wave-ending rule, the late increase cannot be attributed unambiguously to an increase in error variance over time. Overall, this pattern is interpreted as a moderate late-period departure from the constant-$\sigma$ approximation, rather than evidence of severe heteroscedasticity.

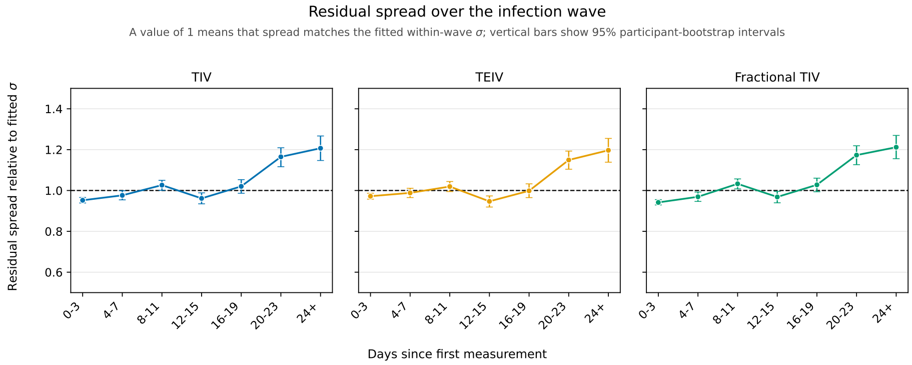


Figure 3: Residual spread by time since the first measurement in each fitted wave. Points show $R_b$, the dashed line denotes the reference value of 1, and vertical bars show central 95% participant-bootstrap intervals.

### 1.4 Short-lag residual association

The working observation model assumes that standardized errors are independent across observation times. Independence implies

$$\mathrm{Cov}(Z_i, Z_j) = 0 \qquad (i \neq j).$$

Because $E(Z_i) = 0$, this also implies $E(Z_i Z_j) = 0$. Rather than attempting a complete test of independence, we examined this necessary implication for within-wave time lags of one to seven days.

For a lag of $d$ days, we calculated

$$M_d = \frac{1}{N_d} \sum_{\substack{i,j \text{ in the same wave} \\ t_j - t_i = d}} \widetilde{Z}_i \widetilde{Z}_j,$$

where $N_d$ is the number of observation pairs separated by $d$ days and $\widetilde{Z}_i$ is the censoring-aware standardized residual defined above. With residual means close to zero, $M_d$ provides a simple covariance-like summary. For the latent errors under the working model, the reference value is zero.

For all three models, the mean residual products were negative at lags of one to five days, reaching approximately $-0.16$ to $-0.18$ at lags of two and three days. They were close to zero at six days and slightly positive at seven days. The strongest negative products were moderate in magnitude and had largely disappeared by six days. Overall, the diagnostic shows some short-range residual association, but no large or persistent departure that would make temporal independence unreasonable as a working approximation.

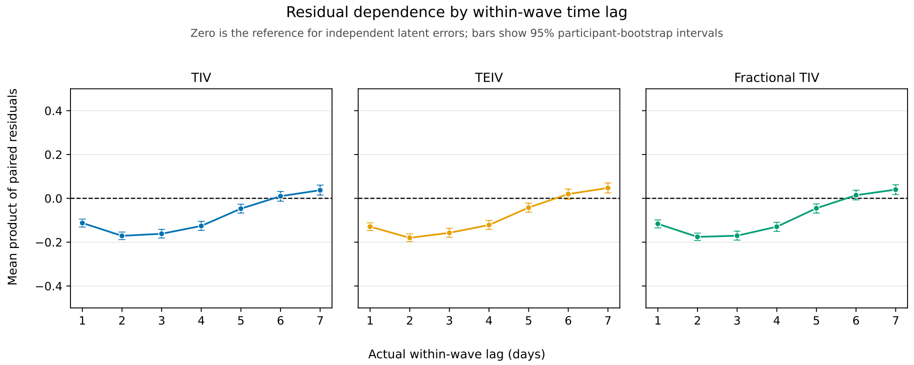


Figure 4: Mean products of censoring-aware standardized residual pairs by within-wave time lag. Points show $M_d$, and the dashed line denotes the zero reference for independent latent errors. Vertical bars show central 95% participant-bootstrap intervals with the fitted models held fixed.

### 1.5 Conclusion

The pooled distribution of the censoring-aware standardized residuals showed no gross departure from the standard normal reference, and the diagnostics showed no substantial time-dependent mean bias. Moderate late-period excess spread and short-range negative residual association were nevertheless present. Overall, the diagnostics support retaining the censored-Gaussian likelihood as a reasonable working approximation.
